# Emergence of a novel hypervirulent extensively drug-resistant ST383 *Klebsiella pneumoniae* lineage carrying ICEKp5 in Lebanon

**DOI:** 10.64898/2026.03.17.712279

**Authors:** Martine Abboud, Tia Clara Chaaya, Yara Daccache, Nour El Alam, Theresa Gerges, Lea Haddad, Leon Kassabian, Jame Tannous, Yara Ghanem, Judy Nabbout, Kevin Chaar, Toni Nmeir, Anthony Haddad, Charbel Al Khoury, George F. Araj, Sima Tokajian

## Abstract

*Klebsiella pneumoniae* ST383 has emerged as a high-risk clone, characterized by carbapenem resistance and increasing detection of hypervirulence determinants. We describe a novel ST383 lineage in Lebanon, defined by the acquisition of ICEKp5, which carries the yersiniabactin locus. Three ST383 *K. pneumoniae* clinical isolates (LBN_CAKp91, LBN_CTKp3, LBN_CTKp11) recovered from a Lebanese medical center were subjected to whole-genome sequencing. Comparative genomic analysis included regional ST383 strains and previously characterized Lebanese isolates. The study isolates formed a tight, monophyletic cluster (3-9 SNPs) that is phylogenetically distinct from the previously reported Lebanese ST383 clone (>164 SNPs) and grouped most closely to an Egyptian ST383 strain (59-65 SNPs). All three isolates carried ICEKp5 with yersiniabactin lineage *ybt14*, a feature absent in the earlier Lebanese ST383 clone. The isolates were the only ST383 strains to harbor the full spectrum of hypervirulence determinants to date, including capsule regulators (*rmpA, rmpA2*), aerobactin (*iucABCD, iutA*), yersiniabactin, and the hypervirulence biomarker *peg-344*. All isolates carried dual carbapenemases (*bla*_OXA-48_ and *bla*_NDM-5_) in addition to *bla*_CTX-M-15_ and *bla*_CTX-M-14b_. The genetic environments of *bla*_OXA-48_ and *bla*_NDM-5_ were highly conserved across geographically diverse ST383 isolates, indicating common plasmid origins. This study documents the emergence of a novel hypervirulent extensively drug-resistant (XDR) ST383 *K. pneumoniae* lineage in Lebanon. The acquisition of ICEKp5, combined with plasmid-borne hypervirulence and resistance determinants, reveals the concerning convergence of hypervirulence and XDR. Enhanced surveillance and infection control measures are urgently needed to monitor this emerging high-risk clone.

## Introduction

A major public health threat has emerged through the convergence of hypervirulence and carbapenem resistance in *Klebsiella pneumoniae*, giving rise to carbapenem-resistant hypervirulent *K. pneumoniae* (CR-hvKp) strains (1). Convergence occurs either when CRKp acquire hypervirulence loci, including potent siderophore systems, and regulators of the mucoid phenotype (*rmpA* and *rmpA2*), or when hvKp acquire carbapenemase genes, frequently through large hybrid plasmids that form through recombination and fusion of distinct backbones (e.g., IncHI1B and IncFIB), enabling simultaneous multidrug resistance and heightened virulence (2, 3).

Multiple high-risk *K. pneumoniae* STs have emerged globally, including ST383 (Mediterranean/African) (4–6), ST11 (Asia), ST23 (worldwide), ST101 (Europe-Asia), ST258 (Americas/Europe) (7), and ST147, ST307, ST15 (8, 9). These clones exhibit distinct geographic distributions, resistance mechanisms, and virulence profiles.

ST383 has emerged as a prominent lineage across Lebanon, Egypt, and Qatar, although with variable prevalence. In Egypt, ST383 represented 41% of colistin-resistant ICU isolates in one cohort (10) and 14.7% of carbapenem-resistant bloodstream isolates in another (11). In Qatar, ST383 was less frequent overall (12), but most affected patients had recently traveled to Egypt, suggesting cross-border dissemination within the Eastern Mediterranean region (EMR).

In Lebanon, ST383 accounted for 76% of carbapenem- and ceftazidime-avibactam-resistant isolates at a tertiary care center (6). Among these, five ST383 isolates carried NDM-5 and OXA-48, highlighting the establishment of this clone in the country. However, the hypervirulence potential of Lebanese ST383 strains has not been fully characterized.

This study reports the emergence of a phylogenetically distinct ST383 *K. pneumoniae* lineage in Lebanon, characterized by ICEKp5, which carries the yersiniabactin virulence locus. Through comprehensive genomic analysis, we aimed to: characterize the virulence and resistance profiles of three clinical isolates, define their phylogenetic relationships with regional ST383 strains, and compare their genetic features with those of the previously reported Lebanese ST383 clone, which lacked ICEKp5 and its associated hypervirulence determinants.

## Methods

### Ethical consideration

Ethical approval was not required for this study since the isolates were collected as part of routine clinical care. Patient-identifying information was anonymized prior to the start of the work.

### Bacterial strains collection and identification

Three non-duplicate clinical *K. pneumoniae* isolates were collected from the Clinical Microbiology Laboratory at the American University of Beirut Medical Center in Lebanon. Two isolates, designated LBN_CTKp3 and LBN_CTKp11, were collected in January 2025, whereas the third, designated as LBN_CAKp91, was collected in May 2025. The isolates were identified using matrix-assisted laser desorption ionization-time-of-flight mass spectrometry (MALDI-TOF MS) with the MALDI Biotyper software (Bruker Daltonics, Bremen, Germany). Carbapenem resistance was determined using the E-test method (AB BIODISK, Solna, Sweden) by testing the minimal inhibitory concentrations (MICs) of ertapenem, imipenem, and meropenem, with results interpreted according to the Clinical Laboratory Standards Institute guidelines(13). The phenotypic detection of carbapenemase types was performed using the NG-Test CARBA-5 (Biotech SAS, Guipry, France), following the manufacturer’s instructions. For further analysis, the isolates were cultured on tryptic soy agar, then preserved in 50% glycerol at -20°C and -80°C.

### Antimicrobial susceptibility testing

The susceptibility of each isolate was determined using the Kirby-Bauer disk diffusion method. A saline suspension of each culture was plated on Mueller-Hinton agar, and the following antibiotic discs were positioned over the inoculum: Ciprofloxacin (5µg), levofloxacin (5µg), ofloxacin (5µg), norfloxacin (10µg), trimethoprim–sulfamethoxazole (1.25/23.75µg), fosfomycin (200µg), nitrofurantoin (100µg), amikacin (30µg), amoxicillin–clavulanic acid (20/10µg), piperacillin–tazobactam (100/10µg), ertapenem (10µg), imipenem (10µg), meropenem (10µg), cefepime (30µg), cefotaxime (30µg), cefixime (5µg), tetracycline (30µg), tigecycline (15µg), gentamicin (10µg), colistin (10µg), cefoxitin (30µg), cefamandole (30µg), cefuroxime (30µg), and aztreonam (30µg) (Bio-Rad, Hercules, CA, USA). After incubation at 37 °C for 24 h, the results were interpreted according to CLSI (13). Isolates were classified as multidrug-resistant (MDR) if resistant to at least one antimicrobial agent in three or more classes, extensively drug-resistant (XDR) if resistant to at least one agent in all but two or fewer categories, and pan-drug resistant (PDR) if non-susceptible to all agents in all antimicrobial categories (14).

### DNA extraction and whole-genome sequencing

Genomic DNA was extracted from overnight cultures of the isolates using the Sigma-Aldrich DNA extraction kit (Sigma-Aldrich; St. Louis, USA) following the manufacturer’s instructions. Genomic libraries were constructed using the Nextera XT DNA Library Preparation Kit (Illumina). Libraries were sequenced on an Illumina MiSeq using a paired-end 500-cycle protocol with a read length of 2×250bp.

### PCR-based replicon typing

Characterization of plasmid incompatibility (Inc) groups was performed using the PBRT 2.0 kit (DIATHEVA, Fano, Italy) according to the manufacturer’s instructions to amplify 30 replicons found among the *Enterobacterales*. The resulting products were visualized on a 2.5% agarose gel stained with ethidium bromide.

### Genome assembly, annotation, and analysis

FastQC v0.11.8 was used to assess the quality of Illumina raw reads (15). Trimmomatic v0.39 was used for adapter removal and trimming (16). De novo genome assembly was done using SPAdes Genome Assembler v3.15.3 (17).

The draft genomes were annotated using the Bakta v1.11.4 pipeline with the v6.0 light database (18). Kleborate v3.2.4 was used for species identification, multilocus sequence typing, detection of virulence markers, and identification of integrative and conjugative elements (ICE) of *K. pneumoniae* (19). Virulence-associated genes were additionally cross-referenced using ABRicate v1.2.0 (20) against the Virulence Factor Database (VFDB) (https://www.mgc.ac.cn/VFs/). The capsule and lipopolysaccharide types were determined using Kaptive v3 (21). Antimicrobial resistance genes and chromosomal mutations were identified using Kleborate (19). Furthermore, acquired antibiotic, biocide, and heavy-metal-resistance genes were identified using AMRFinderPlus v4.2.5 with the “plus” database v2025-12-03.1 (22). Abricate v1.2.0 was used to detect fitness-associated genes (20).

Plasmid replicons were identified using PlasmidFinder v2.1.6 (https://cge.cbs.dtu.dk/services/PlasmidFinder/), and further validated using the MOB-typer module of MOB-suite v3.1.9 (23).

### Phylogenetic tree construction and visualization

Core SNPs were identified using Snippy v4.6 by aligning the reads from each genome (https://github.com/tseemann/snippy, accessed on 25 January 2021). High-quality SNPs and indels were annotated using SnpEff v5.1 after removal of low-confidence alleles with a consensus base quality <20 and a read depth <5 or a heterozygous base calls (24). The parsimony tree was constructed from consensus core genome sequences generated by Snippy and kSNP3 v3.0 (25). The final output was visualized using iTOL (26).

### Graphical representation and visualization

The hypervirulence gene heatmap (Figure 3B) was generated using Python v3.12 with the matplotlib and pandas libraries. Gene presence values were derived from Kleborate v3.2.4 output and cross-referenced with ABRicate v1.2.0 against the VFDB and AMRFinderPlus v4.2.5. All other graphs were generated in R v.3.6.1 using ggplot2, ggrepel, pheatmap, and ComplexHeatmap. Pairwise genomic alignments were visualized using clinker, with gene links displayed for pairwise amino acid identities exceeding 65%.

### Data availability

The draft genomes have been deposited at the National Center for Biotechnology Information (NCBI) database. The accession numbers for these isolates, and for those retrieved from NCBI and included in the analysis, are listed in Supplementary S1.

## Results

### SNP-based phylogenetic analysis

SNP-based phylogenetic reconstruction of ST383 *K. pneumoniae* isolates revealed distinct evolutionary relationships associated with geographic origins (Figure 1). Our study isolates (LBN_CAKp91, LBN_CTKp3, and LBN_CTKp11) formed a well-supported monophyletic clade, with low pairwise SNP distances (3-9 SNPs), and grouped most closely with EGY_DAPDKI01.1. LBN_CTKp3 exhibited only 59 SNPs differences from EGY_DAPDKI01.1, while LBN_CAKp91 and LBN_CTKp11 showed 60 and 65 SNPs, respectively, from this Egyptian strain.

**Figure 1:**
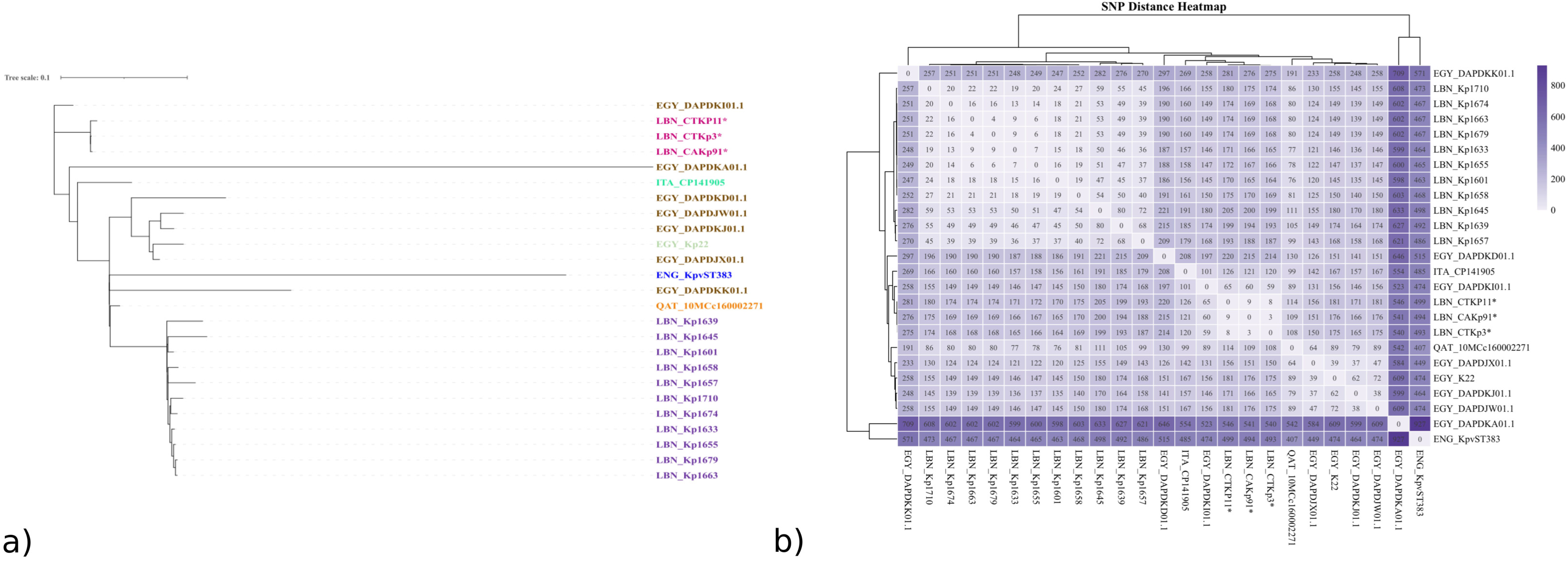
SNP-based phylogenetic analysis of ST383 *Klebsiella pneumoniae* isolates. a. SNP-based phylogenetic tree representing the evolutionary relationships among ST383 *K. pneumoniae* isolates from different geographic origins. Branch lengths are scaled to represent the number of substitutions/site (scale bar = 0.1). Isolates are color-coded by country of origin. The tree was generated using Snippy v. 4.6 and visualized through iTOL b. SNP distance heatmap displaying pairwise genetic divergence between isolates. Color intensity corresponds to the total SNP count between strains, with numerical values shown in each cell. Lower values (lighter colors) indicate closer genetic relationships. Isolates marked with an asterisk (*) represent the current study, while unmarked strains were obtained from NCBI. The heatmap was generated using R v. 3.6.1

The next closest were the isolates from Qatar (108–114), Italy (120–126), and another from Egypt (150–156). In contrast, a substantial divergence was observed from the Sobh et al. (2024) Lebanese isolates (range: 164-220 SNPs). The EGY_K22 isolate, previously reported to carry similar virulence determinants including the aerobactin locus and IncHI1B/IncFIB virulence plasmid, showed SNP distances of 175-181 from our isolates. These phylogenomic data reveal a distinct ST383 introduction into Lebanon, closer to Egyptian strains than previous local endemic lineages.

### Comparative analysis of antibiotic resistance genes and plasmid replicons

The three isolates from this study (LBN_CTKp3, LBN_CAKp91, and LBN_CTKp11) were extensively drug-resistant (XDR), exhibiting phenotypic resistance to all tested antibiotics except colistin; LBN_CAKp91 was additionally susceptible to tigecycline. Comparative analysis of antibiotic resistance genes and plasmid replicons revealed distinct resistome profiles among ST383 *K. pneumoniae* isolates (Figure 2a).

**Figure 2:**
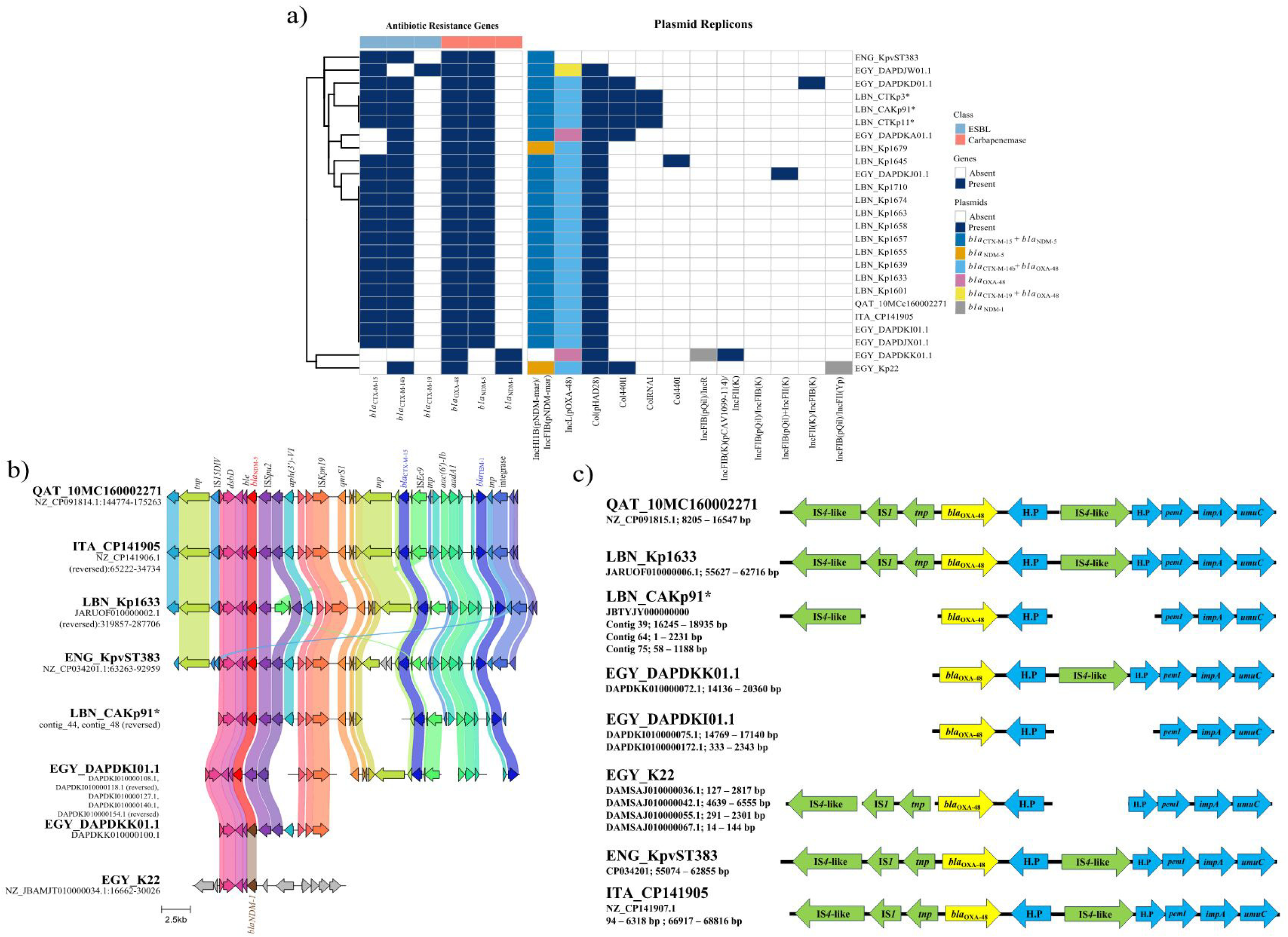
Distribution of antimicrobial resistance genes, plasmid replicons, and carbapenemase genetic contexts in ST383 *Klebsiella pneumoniae* isolates. a. Heatmap of antibiotic resistance genes and plasmid replicons in ST383 *K. pneumoniae* isolates. Heatmap displaying the presence of antibiotic resistance genes (left panel) and plasmid replicons (right panel) across ST383 *K. pneumoniae* isolates. Resistance genes are grouped by class (color-coded bar at top): ESBL genes (*bla*_CTX-M-15_, *bla*_CTX-M-14b_, *bla*_CTX-M-19_) and carbapenemase genes (*bla*_OXA-48_, *bla*_NDM-5_, *bla*_NDM-1_). Dark blue indicates gene/plasmid presence, while white indicates absence. Plasmid replicons are color-coded to indicate associated resistance genes. Isolates are hierarchically clustered based on resistance and plasmid profiles (dendrogram, left). Isolates from the current study are marked with asterisks (*). The heatmap was generated using R v. 3.6.1. b. Comparative analysis of the *bla*_NDM_ genetic environment in ST383 *K. pneumoniae* isolates. Linear comparison of the genetic regions flanking *bla*_NDM_ across ST383 *K. pneumoniae* isolates from different geographic origins. Genes are represented as arrows indicating transcriptional direction and are color-coded: *bla*_NDM_, transposases and insertion sequences (*tnp*, IS*15*DIV, IS*Spu2*, IS*Kpn19*, IS*Ec9*), bleomycin resistance (*ble*), aminoglycoside resistance genes (*aph(3’)*, *aac(6’)-Ib*, *aadA1*), quinolone resistance (*qnrS1*), and *bla*_TEM-1._ Colored ribbons connect homologous regions between isolates, illustrating the conserved synteny. Scale bar represents 2.5 kb. Isolate names and contig coordinates are shown on the left; the study isolate is marked with an asterisk (*). Structural variations in some isolates reflect fragmentation of draft genome assemblies across multiple contigs. c. Comparative analysis of the *bla*_OXA-48_ genetic environment in ST383 *K. pneumoniae* isolates. Linear representation of the genetic regions flanking the *bla*_OXA-48_ across ST383 *K. pneumoniae* isolates from different geographic origins. Genes are represented as arrows indicating transcriptional direction and are color-coded: the *bla*_OXA-48_ (yellow), insertion sequences and transposases (IS*4*-like, IS*1*, *tnp*; green), hypothetical proteins (H.P; green), and other genes (*pemI*, *impA*, *umuC*; blue). Isolate names and contig coordinates are shown on the left. All isolates share a conserved genetic environment consisting of IS*4*-like–IS*1*–*tnp* upstream of the *bla*_OXA-48_, followed by a hypothetical protein, IS*4*-like, and the *pemI*-*impA*-*umuC* gene cluster downstream. Incomplete structures in some isolates (LBN_CAKp91, EGY_DAPDKK01.1, EGY_DAPDKI01.1, EGY_K22) reflect the fragmented nature of draft genome assemblies across multiple contigs rather than true genetic variation.

The three study isolates shared an identical resistome with the previously reported Lebanese ST383 isolates, despite their phylogenetic divergence. This shared resistome comprised the ESBL genes *bla*_CTX-M-15_ and *bla*_CTX-M-14b_ alongside the carbapenemase genes *bla*_NDM-5_ and *bla*_OXA-48_, distributed across the same plasmid backbones: an IncL plasmid carrying *bla*_CTX-M-14b_ and *bla*_OXA-48_, and a hybrid IncHI1B(pNDM-MAR)/IncFIB(pNDM-Mar) plasmid carrying *bla*_CTX-M-15_ and *bla*_NDM-5_. These genes were conserved across all ST383 isolates analyzed, with two exceptions: ENG_KpvST383 lacked *bla*_OXA-48_, and EGY_K22 and EGY_DAPDKK01.1 harbored *bla*_NDM-1_ instead of *bla*_NDM-5_.

Among phylogenetically related strains, EGY_DAPDKA01.1 and ITA_CP141905 shared a similar resistome but differed in their plasmid content: Col440II plasmids were absent in EGY_DAPDKA01.1, and ColRNAI was detected exclusively in our study isolates. In contrast, the hybrid IncHI1B(pNDM-MAR)/IncFIB(pNDM-MAR) plasmid, a hallmark of ST383, was present in all strains. The Egyptian isolates displayed more variable ESBL combinations: EGY_DAPDJW01.1 carried *bla*_CTX-M-19_ and *bla*_OXA-48_ on an IncL plasmid, while EGY_K22 harbored *bla*_CTX-M-14b_ and *bla*_OXA-48_ on IncL together with *bla*_NDM-1_. The co-occurrence of *bla*_NDM-5_, *bla*_OXA-48_, *bla*_CTX-M-15_, and *bla*_CTX-M-14b_ constitutes the predominant resistance profile among ST383 isolates in the EMR.

### Genetic environment of *bla*_NDM_

Two *bla*_NDM_ variants were identified across eight representative ST383 isolates from five countries: *bla*_NDM-1_ in EGY_DAPDKK01.1 and EGY_K22, and *bla*_NDM-5_ in the six remaining isolates (Figure 2b).

In QAT_10MC160002271, ITA_CP141905, LBN_Kp1633, and ENG_KpvST383, *bla*NDM-5 is embedded within a conserved IS*15DIV*-*dsbD-ble-bla*_NDM-5_-IS*Spu2* segment, maintained in the same gene order across these complete assemblies and consistent with an insertion sequence–mediated mobile unit. In LBN_CAKp91 and EGY_DAPDKI01.1, contig fragmentation precluded assessment of the upstream IS*15DIV*-containing region; however, the *dsbD*–*ble*–*bla*_NDM_–IS*Spu2* core was identifiable in all three, suggesting conservation of this cassette.

Downstream of IS*Spu2*, QAT_10MC160002271, ITA_CP141905, LBN_Kp1633, and ENG_KpvST383 carried an identical MDR region (Figure 6). LBN_CAKp91 and EGY_DAPDKI01.1 showed the same downstream gene arrangement, although *aph(3’)-VI* was not detected within the aligned segment of EGY_DAPDKI01.1.

EGY_DAPDKK01.1 shared the conserved upstream of the *bla*_NDM-1_ cassette, yet downstream of IS*Spu2*, it retained only *aph(3’)-VI* and IS*Kpn19*. This pattern supports a conserved IS-bounded *bla*_NDM_ module acquired on a common backbone, followed by lineage-specific assembly or loss of the extended MDR region.

EGY_K22 showed a clearly distinct organization. Upstream of *bla*_NDM-1_, only *dsbD* and *ble* were present, with no IS*15DIV* or IS*Spu2* detected. In addition, none of the resistance genes found in the conserved downstream array of the other isolates were identified within the aligned segment.

### Genetic environment of *bla*_OXA-48_

Analysis of the genetic environment surrounding *bla*_OXA-48_ revealed a conserved arrangement among ST383 *K. pneumoniae* isolates (Figure 2c). It comprised IS*4*-like and IS*1* insertion sequences upstream of a transposase gene (*tnp*) preceding *bla*_OXA-48_, followed by a hypothetical protein (HP), an additional IS*4*-like element, and the conserved *pemI*-*impA*-*umuC* cluster downstream. The variations in the representation of flanking elements in some isolates (e.g., LBN_CAKp91, EGY_DAPDKK01.1, EGY_DAPDKI01.1) are attributable to the draft nature of their genome assemblies, where *bla*_OXA-48_ and its flanking regions are distributed across multiple contigs rather than representing true structural differences.

### Comparative analysis of hypervirulence gene profiles

Comparative analysis of hypervirulence-associated genes revealed distinct virulence profiles among ST383 isolates (Figure 3b). Our three isolates were the only strains to harbor the complete spectrum of hypervirulence determinants, including the capsule regulators *rmpA* and *rmpA2*, complete aerobactin siderophore (*iucABCD* and *iutA*) and yersiniabactin siderophore (*ybtAEPQSTUX*) loci, and the hypervirulence biomarker *peg-344*, all with high sequence identity. In contrast, all other ST383 isolates analyzed exhibited incomplete hypervirulence gene profiles. The Egyptian strain EGY_DAPDKI01.1 lacked *rmpA* and *rmpA2*. The Lebanese ST383 isolates reported by Sobh et al. (2024) contained the aerobactin (*iuc*) locus in all isolates except one. Low-identity matches to the capsule regulators *rmpA* and/or *rmpA2* were detected in a subset of isolates, whereas the *yersiniabactin (ybt)* locus and *peg-344* were absent from all genomes. The current study isolates represent the most hypervirulent sublineage of ST383 identified to date, with a virulence score of 4 (Figure 3b), uniquely characterized by the convergence of all major virulence determinants.

**Figure 3:**
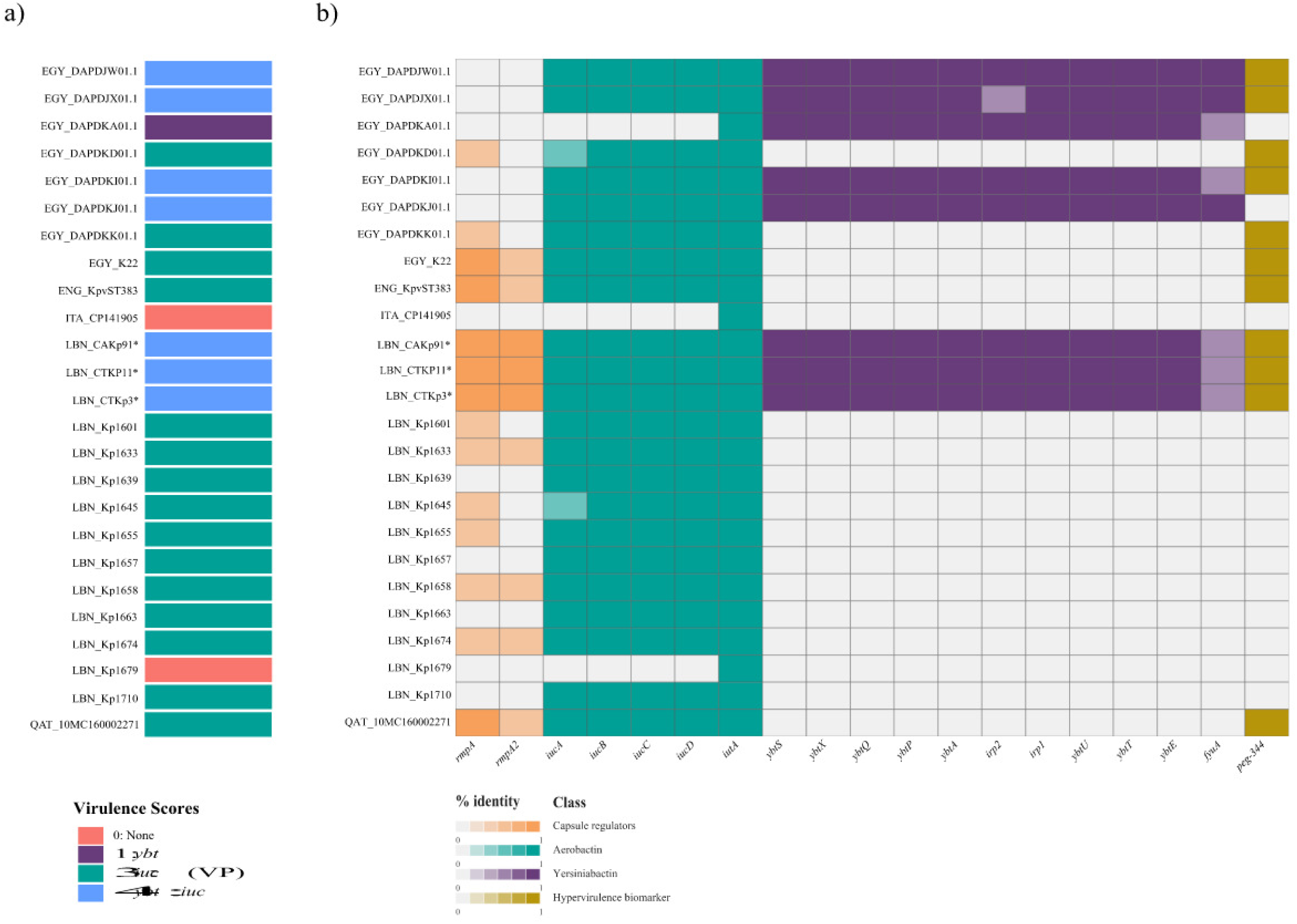
Hypervirulence gene profiles across ST383 *klebsiella. pneumoniae* isolates. a. Virulence score profiles across ST383 *K. pneumoniae* isolates. Data shown summarizes Kleborate results; isolates are shown as a single-column heatmap. Different colors indicate virulence score categories based on the presence of key virulence loci: 0 (none), 1 (yersiniabactin; *ybt*), 3 (aerobactin: *iuc*: VP: virulence plasmid), 4 (*ybt* + *iuc*: VP). b. Heatmap displaying the presence and percent nucleotide identity of hypervirulence-associated genes. Genes are grouped by functional class: capsule regulators (*rmpA, rmpA2*), aerobactin siderophore system (iucA, iucB, iucC, iucD, iutA), yersiniabactin siderophore system (*ybtS, ybtX, ybtQ, ybtP, ybtA, irp2, irp1, ybtU, ybtT, ybtE, fyuA*), and hypervirulence biomarker (*peg-344*). Color intensity represents percent nucleotide identity (0–1 scale), with full color indicating high identity and white indicating gene absence. Study isolates are marked with asterisks (*).

### Structural organization of the virulence loci on IncHI1B/IncFIB hybrid virulence plasmid

Due to identical genomic structures and minimal pairwise distances (3-9 SNPs), LBN_CAKp91 was selected as the representative study isolate in the comparative alignment; all observations apply equally to LBN_CTKp3 and LBN_CTKp11.

The virulence region of the hybrid plasmid across seven representative ST383 isolates from Lebanon, Egypt, England, and Qatar showed a conserved central backbone comprising the aerobactin (*iucABCD-iutA*) and the tellurite resistance (*terZABCDW*) operons (Figure 4).

**Figure 4:**
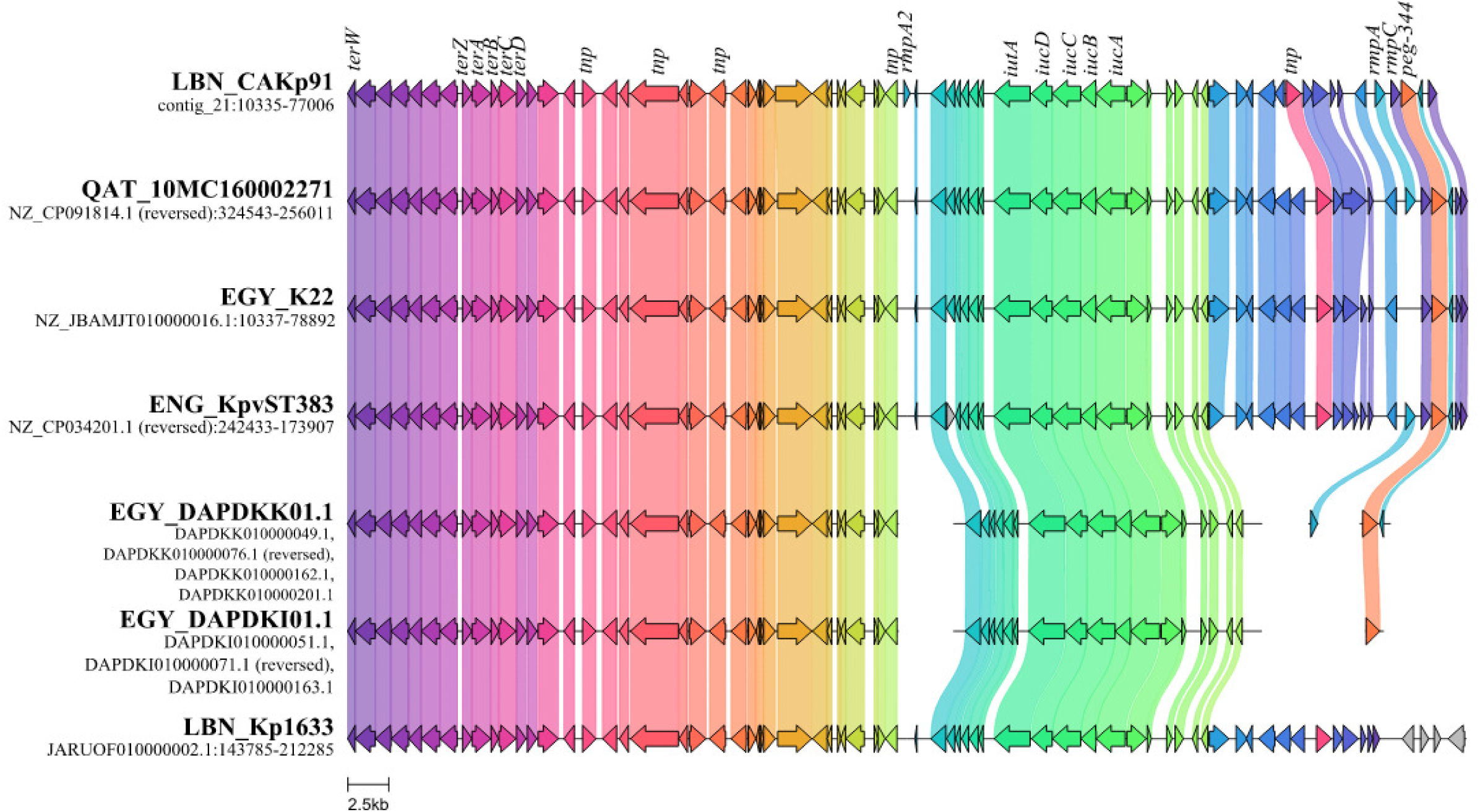
Structural organization of the virulence-associated plasmid region across representative ST383 *Klebsiella pneumoniae* isolates. Seven isolates spanning Lebanon (LBN), Egypt (EGY), England (ENG), and Qatar (QAT) are compared across a region encompassing the tellurite resistance operon (*terZABCDW*), aerobactin biosynthesis and uptake locus (*iucABCD*/*iutA*), and the hypermucoviscosity-associated cassette (*rmpA*, *rmpC*, *rmpA2*, *peg-344*). Contig fragmentation in Egyptian draft assemblies reflects limitations of short-read sequencing and should be considered when interpreting apparent gene absence in the distal flanking regions. Asterisks denote study isolates.

Variation was restricted to the flanking *rmpA*, *rmpC*, and *peg-344*. LBN_CAKp91, ENG_KpvST383, and QAT_10MC160002271 retained the complete cassette, whereas EGY_K22 and EGY_DAPDKK01.1 lacked *rmpA* and *rmpC*, respectively. EGY_DAPDKI01.1 retained only *peg-344*, and LBN_Kp1633 lacked all three elements.

As the Egyptian genomes are draft assemblies, the absence in this region should not be interpreted as a confirmed gene loss. Lastly, LBN_CAKp91 displayed the most complete configuration, carrying *rmpA* and *rmpA2*, though truncated.

### ICEKp comparative genomic structure analysis

Comparative analysis of ICEKp elements demonstrated a conserved core organization across all eight ICE-positive ST383 isolates, independent of geographic origin or ICEKp type (Figure 5). All elements were integrated at the chromosomal tRNA-Asn site and shared the same structural framework: an integrase gene at the left boundary, the complete yersiniabactin biosynthesis and transport locus (*ybtS, ybtX, ybtQ, ybtP, ybtA, irp2, irp1, ybtU, ybtT, ybtE*, and *fyuA*), a type IV secretion system (T4SS) region at the right boundary (*virB1, virB2, virB5, virB8, virB9, virD4*), and *mobC*. The yersiniabactin locus was intact in all isolates, with no internal disruptions in the recovered regions.

**Figure 5:**
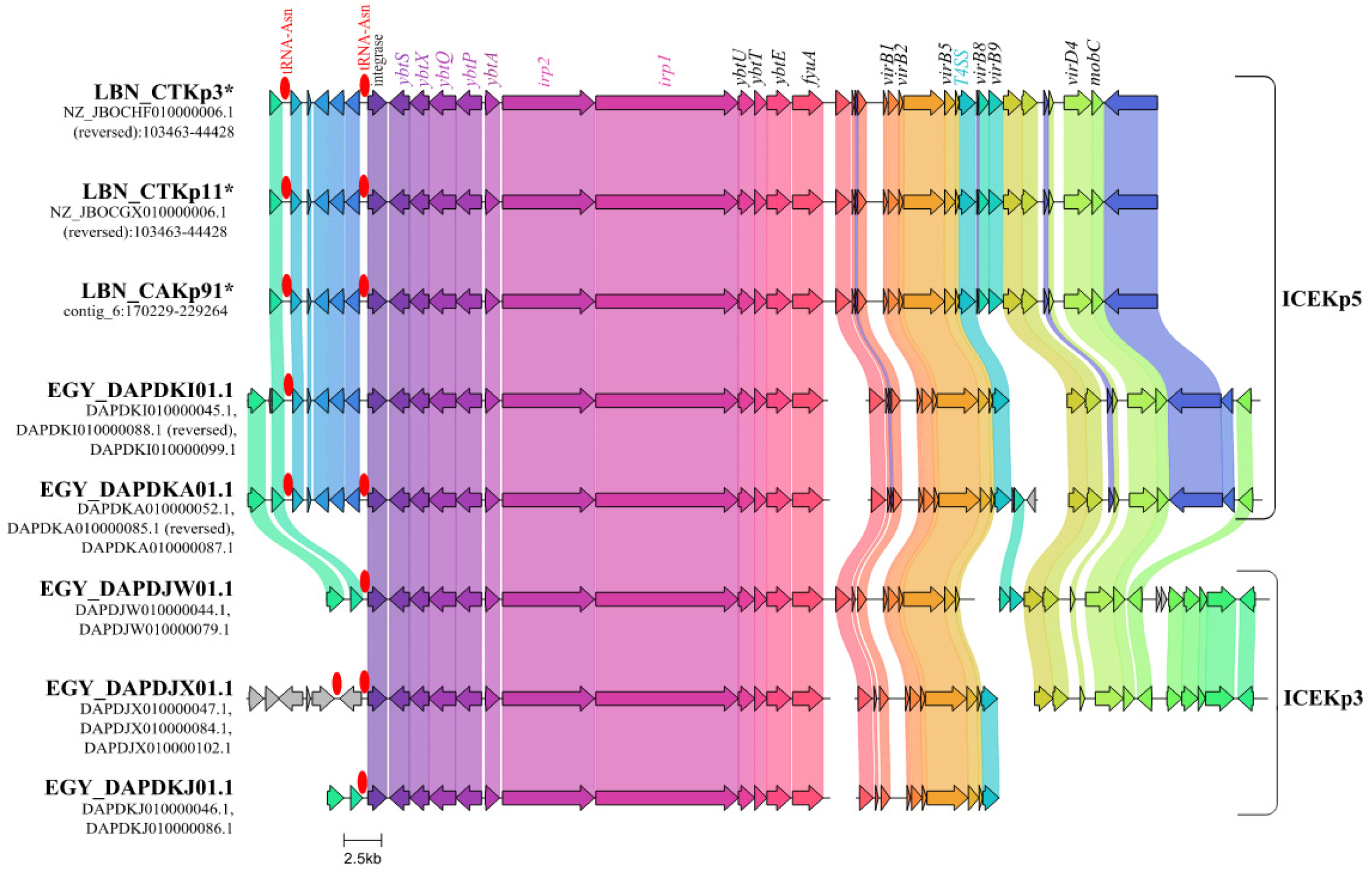
Linear comparison of ICEKp element structure across ST383 *Klebsiella pneumoniae* isolates. Gene blocks are represented as arrows indicating transcriptional direction and are colored by sequence identity, with colored ribbons connecting homologous regions between isolates. Red ovals denote the chromosomal tRNA-Asn integration sites characteristic of ICEKp elements. The conserved ICE architecture includes: an integrase gene at the left boundary; the complete yersiniabactin biosynthesis and transport locus (*ybtS*, *ybtX*, *ybtQ*, *ybtP*, *ybtA*), *irp2* and *irp1*, and *ybtU*, *ybtT*, *ybtE*, *fyuA*; and the Type IV Secretion System (T4SS) conjugation machinery including *virB1*, *virB2*, *virB5*, T4SS components, *virB8*, *virB9*, and *virD4*, *mobC* at the right boundary. Scale bar represents 2.5 kb. Isolates are bracketed by ICEKp type as assigned by Kleborate: ICEKp5 (study isolates LBN_CTKp3*, LBN_CTKp11*, LBN_CAKp91*, and Egyptian isolates EGY_DAPDKI01.1, EGY_DAPDKA01.1) and ICEKp3 (EGY_DAPDJX01.1, EGY_DAPDKJ01.1, EGY_DAPDJW01.1). Asterisks (*) denote study isolates. Contig breaks in Egyptian draft assemblies reflect short-read fragmentation at repetitive ICE boundaries; synteny is preserved across assembled segments.

T4SS genes *virB1* and *virB2* were detected in all genomes, supporting retention of conjugative potential of the yersiniabactin locus. Contig breaks in the Egyptian draft assemblies were consistent with short-read fragmentation at repetitive ICE boundaries rather than true structural rearrangement, as gene order remained conserved across aligned segments.

Despite this shared backbone, two distinct ICEKp configurations were observed, matching Kleborate assignments. The three study isolates, LBN_CTKp3, LBN_CTKp11, and LBN_CAKp91, carried ICEKp5 with *ybt14* locus. Egyptian isolates, EGY_DAPDKI01.1 and EGY_DAPDKA01.1 also carried ICEKp5, whereas EGY_DAPDJW01.1, EGY_DAPDJX01.1, and EGY_DAPDKJ01.1 carried ICEKp3 with *ybt9*. The ICEKp3-positive isolates exhibited differences in flanking gene content compared with ICEKp5. EGY_DAPDJW01.1 and EGY_DAPDKJ01.1 were assigned YbST 183, while EGY_DAPDJX01.1 was assigned YbST 173-2LV due to two novel variants in *irp2* and *irp1* within the ICEKp3 backbone. All five ICEKp5-positive isolates shared YbST 151-1LV, a single variant of YbST 151, supporting a single ICEKp5 acquisition event from a common ancestor.

### Convergent evolution of ST383 in Lebanon toward an hv-XDR phenotype

The three study isolates, CTKp3, CTKp11, and CAKp91, represent a newly emerging ST383 lineage converging hypervirulence and extensive drug resistance through stepwise acquisition of mobile genetic elements. ST383, an independent non-CG258 lineage, has progressed over the past decade from an OXA-48-producing ESBL clone first described in the United Kingdom in 2019 to a globally disseminated high-risk clone across the EMR. These isolates carry an ICEKp5 (∼75–80 kb) integrated at the tRNA-Asn site, encoding the yersiniabactin locus *ybt14* and a functional T4SS, alongside an IncL pOXA-48 plasmid and an IncFIB/IncHI1B hybrid plasmid encoding *bla*_NDM-5_, *bla*_CTX-M-15_, the aerobactin operon (*iucABCD*, *iutA*), and hypermucoidy regulators *rmpA* and *rmpA2*. This genomic configuration yields a Kleborate virulence score of 4, classifying these isolates as convergent hypervirulent

## Discussion

*K. pneumoniae* ST383 has emerged as a high-risk lineage increasingly associated with the convergence of MDR and hypervirulence across the EMR and beyond (6, 10, 12, 27). In Lebanon, ST383 was previously recognized as a carbapenem-resistant hospital-adapted clone carrying *bla*_NDM-5_ and *bla*_OXA-48_ on IncFIB-IncHI1B and IncL plasmids, respectively (6). In that cohort, 26 of 34 *K. pneumoniae* isolates recovered between 2019 and 2021 belonged to ST383 and differed by only 4-80 SNPs, consistent with local clonal expansion. However, it lacked evidence of hypervirulence-associated elements such as ICEKp5. This study reports the emergence of a phylogenetically distinct ST383 *K. pneumoniae* lineage in Lebanon, defined by the acquisition of ICEKp5, which carries the yersiniabactin virulence locus, and harbors the full complement of hypervirulence determinants and dual carbapenemase genes. These findings highlight the ongoing convergence of hypervirulence and XDR within this high-risk clone.

The three *K. pneumoniae* ST383 isolates characterized in this study are distinct (164-220 SNPs) from the previous Lebanese ST383 isolates (6), indicating an independent introduction of ST383 into Lebanon. The study isolates clustered closely with Egyptian ST383 strains, particularly EGY_DAPDKI01.1. LBN_CTKp3, LBN_CAKp91, and LBN_CTKp11 exhibited 59, 60, and 65 SNPs, respectively, from this Egyptian strain. The next closest were the isolate from Qatar (108–114), Italy (120–126), and another from Egypt (150–156). This phylogenetic proximity to Egyptian and regional strains suggests a potential epidemiological link, possibly through healthcare-associated travel, patient transfer, or shared transmission networks within the EMR.

A key feature distinguishing the present isolates from those reported by Sobh et al. (2024) is the presence of ICEKp5 carrying the yersiniabactin locus *ybt14* (*ybt* ST151). This element, integrated at the chromosomal tRNA-Asn site, contains the complete yersiniabactin biosynthesis and transport locus and a functional T4SS enabling conjugative transfer. In addition, our isolates harbor a convergent IncHI1B/IncFIB plasmid encoding the full aerobactin locus (*iucABCD* and *iutA*) together with hypermucoidy regulators (*rmpA, rmpA2, rmpC,* and *rmpD*). This combination results in a Kleborate virulence score of 4, classifying them as convergent XDR hypervirulent strains. The co-localization of the tellurite resistance operon *terZABCDW* with the aerobactin locus and hypermucoidy regulators on the same plasmid backbone suggests that tellurite exposure in hospital environments may represent an additional, antibiotic-independent selective pressure favoring plasmid maintenance and contributing to the nosocomial persistence of this convergent clone. The absence of the colibactin (*clb*) and salmochelin (*iro*) loci further defines a specific virulence gene repertoire characteristic of this ST383 sublineage (12).

Among Egyptian isolates, EGY_DAPDKI01.1 and EGY_DAPDKA01.1 carried ICEKp5, whereas EGY_DAPDJW01.1, EGY_DAPDJX01.1, and EGY_DAPDKJ01.1 carried ICEKp3 with *ybt9* (10). All five ICEKp5-positive isolates shared YbST 151-1LV, a single variant of YbST 151, supporting a single ICEKp5 acquisition event from a common ancestor. The conserved ICEKp5 configuration across the three Lebanese study isolates and two Egyptian strains supports close phylogenetic relatedness and defines a distinct sub-lineage separate from the ICEKp3-dominant Egyptian ST383 isolates that have emerged within the EMR. The yersiniabactin siderophore system is important for iron acquisition in nutrient-limited host environments such as the urinary tract and bloodstream (28). Yersiniabactin has been established as a virulence factor for *K. pneumoniae* during pulmonary infection, with yersiniabactin mutants showing attenuated virulence in murine models (28). The acquisition of ICEKp5 by the study isolates represents a significant step toward convergent hypervirulence. This comparative analysis of hypervirulence-associated genes further showed that the study isolates were the only ST383 strains to harbor the complete spectrum of hypervirulence determinants, including *rmpA* and *rmpA2*, the aerobactin siderophore system (*iucABCD* and *iutA*) on the IncHI1B/IncFIB hybrid virulence plasmid, yersiniabactin on ICEKp5, and the hypervirulence biomarker *peg-344*, which is absent from the Sobh et al. (2024) Lebanese ST383 cohort.

ST383 demonstrates a progressive evolutionary trajectory driven by repeated acquisition and reshuffling of major carbapenemase and virulence determinants. Despite their phylogenetic distinctiveness and divergent virulence profiles, they share an identical resistance gene profile. Globally disseminated ST383 strains from Egypt, Lebanon, Qatar, and Italy (12, 27, 29), co-harbored *bla*_OXA-48_ together with either *bla*_NDM-1_ or *bla*_NDM-5_, ESBLs such as *bla*_CTX-M-15_ and *bla*_CTX-M-14_, indicating the incorporation of different NDM variants during the acquisition of the large IncHI1B/IncFIB hybrid resistance and virulence plasmid. Analysis of the genetic environment surrounding *bla*_OXA-48_ revealed a highly conserved structure across all isolates, suggesting a common origin of the *bla*_OXA-48_ -carrying IncL(pOXA-48) plasmid and its stable maintenance within the ST383 lineage. Similarly, the *bla*_NDM_ genetic environment was highly conserved, with most isolates carrying *bla*_NDM-5_. The conserved structure included an MDR module conferring resistance to carbapenems, aminoglycosides, quinolones, and other β-lactams, severely limiting therapeutic options.

In Lebanon and Egypt, sequenced ST383 isolates predominantly carried *bla*_NDM-5_, whereas *bla*_NDM-1_ on related hybrid plasmids was identified in other lineages such as ST11, demonstrating lineage-specific variation and accessory genome plasticity. Earlier European ST383 isolates carried alternative carbapenemases, including *bla*_VIM-4_, *bla*_VIM-19_, and *bla*_KPC-2_, revealing that this clone has repeatedly replaced resistance determinants over time (30). Phylogenetic and plasmid data support a model of sequential acquisition, with ancestral ST383 first acquiring an IncL plasmid bearing *bla*_OXA-48_, followed by integration of a large IncHI1B fusion plasmid carrying *bla*_NDM_ variants and virulence loci, transforming a conventional MDR lineage into a convergent, highly adaptable pathogen. Our findings indicate that ST383 in Lebanon is not solely persisting through local hospital expansion but embraces a distinct introduced lineage. The detection of ICEKp5, absent from previously reported Lebanese ST383 genomes, along with clear phylogenetic separation, supports an independent introduction event rather than local microevolution of the earlier circulating clone. These findings emphasize that ST383 continues to evolve through regional dissemination and stepwise genetic acquisition, highlighting its growing impact on clinical management and infection control across interconnected healthcare systems.

This study documents the emergence of a novel ST383 *K. pneumoniae* lineage in Lebanon, characterized by convergence of XDR and hypervirulence following acquisition of ICEKp5. The three study isolates uniquely harbor all hypervirulence determinants, including *rmpA, rmpA2*, aerobactin, yersiniabactin, and *peg-344*. By also co-harboring *bla*_OXA-48_ and *bla*_NDM-5_, they represent the most extensively characterized hv-XDR ST383 variant reported to date. Phylogenetic analysis confirmed the independent introduction of these isolates, distinct from the previously endemic Lebanese clone and closely related to strains circulating in the region. These findings highlight regional dissemination, emphasizing the need for enhanced genomic surveillance, strengthened infection-control measures, and coordinated regional efforts to limit the spread of this emerging high-risk clone in the EMR.

The study has several limitations. The draft assemblies fragmented some mobile elements, and the lack of epidemiological data prevented definitive identification of the source of introduction.

